# Effects of amendments on remediation of heavy metal contaminated soil and accumulation of Cd and Pb in potato

**DOI:** 10.1101/2025.03.30.646218

**Authors:** Ren He, Ke Liu, Fan Yang, Shiqing Feng, Lili Zhang

**Author notes:** Corresponding author:Ke Liu,Agricultural College of Guizhou University,550025GuiYang,China, /.

## Abstract

With the development of agriculture and industry, a large amount of heavy metals are loaded into the soil, posing a high ecological and public health risk. Many methods have been proposed to repair heavy metal contaminated soil, such as electroremediation, chemical washing, phytoremediation and in situ fixation. In the above methods, the application of passivators to in-situ immobilization of heavy metals is an environmentally friendly and cost-effective choice. The effects of different amendments on the improvement of heavy metal contaminated soil and the accumulation of Cd and Pb in potato plants were investigated by using attapulgite, diatomite and humic acid as materials.The results showed that soil pH, organic matter content and cation exchange capacity increased after the application of amendments. Under the treatment of attapulgite+humic acid, the soil pH, organic matter content and cation exchange capacity were increased by 0.57units, 10.0 mg/kg and 7.28cmol/kg, respectively, compared with the control. The content of soil hydrolyzable nitrogen increased, and the content of available phosphorus decreased. The application of amendments increased soil enzyme activity; effectively reduce the bioavailability of Cd and Pb in soil ; under the treatment of attapulgite+humic acid, the contents of Cd and Pb in potato plants decreased by 85.50%and 55.54%respectively, and the contents of Cd and Pb in potato tubers decreased to 0.08 mg/kg and 1.41 mg/kg respectively. The application of amendments affected the accumulation and transport of heavy metals Cd and Pb in potato plants to a certain extent and promoted the transformation of Cd and Pb from acid-soluble and reduced states to oxidized and residual states.

## Introduction

Heavy metal pollution has become a key issue of global concern due to its potential harmful risks to human health and the environment(KAMRAN, M, et al., 2021; RIAZ, M, et al., 2021)Northwest Guizhou, China is a high geological background area of karst heavy metals. Affected by the historical zinc smelting method, the heavy metal content of atmospheric dust in the region is high, the soil pollution is serious, and there is a high risk of safe production of crops.

Chemical passivation repair regulates the physical and chemical environment of heavy metals in the soil by adding passivation materials to the soil, reducing the bioavailability and mobility of heavy metals in the soil environment, thereby reducing their toxicity to crops^**Error! Reference source not found**.^ to ensure the safe growth of food crops. It was found that attapulgite had a good passivation effect on Cd and Pb ^**Error! Reference source not found**.^. Du Zhimin et al. ^**Error! Reference source not found**.^ showed that attapulgite reduced the content of exchangeable Cu and Cd in soil, and increased soil fertility and enzyme activity. Zhao Tingwei et al^**Error! Reference source not found**.^ concluded that the addition of palygorskite could significantly reduce the content of available Cd in soil. The application of humic acid can promote the transformation of Pb and Cd from weak acid extractable state to residual state, thus effectively reducing the migration ability of Pb and Cd^**Error! Reference source not found**.^. Humic acid can form humic acid-cadmium complex with available cadmium in soil, reduce the content of available cadmium in soil, and inhibit the absorption of cadmium by plants^**Error! Reference source not found**.^.

To solve the problem of excessive heavy metals in farmland soil, it is urgent to screen soil passivator materials. The purpose of this study is to develop a practical new method to improve the growth of potato and the repair effect of metal by optimizing the combination of various passivation agents. The experimental treatments with different levels of passivators and different treatments can improve enzyme activity, promote antioxidants and soil health, and significantly promote growth. The balanced application of passivation agent effectively reduced the concentration of metal and inhibited the absorption of metal by potato plants. Compared with the combined form, the results of the single application of all drugs were statistically similar. In addition, the passivator indirectly enhanced the activity of essential antioxidant enzymes, and the synergistic effect significantly improved the bioaccumulation of Pb and Cd in potato. The key is that the combined use of passivators can improve the growth of potatoes, improve the antioxidant defense system, and improve the quality of potatoes. By reducing heavy metal pollution, passivators can not only improve crop quality and yield.Based on the theory of previous studies, this study used attapulgite, diatomite and humic acid as modifier materials, and selected the soil seriously polluted by heavy metals in northwestern Guizhou for pot experiments. The effects of different modifiers on the improvement of heavy metal contaminated soil and the absorption and accumulation of heavy metals in soil by potato plants were explored by setting up a single modifier and a composite modifier, in order to provide a feasible theoretical reference for the remediation of heavy metal contaminated soil and provide a scientific basis for crop safety production.

## Materials and methods

### Experimental design

The soil at 0-20cm depth (pH of 5.6, soil organic matter of 15.85g / kg, soil heavy metal content of Cd : 6.42mg / kg, Pb : 106.7mg / kg) was collected from the soil surface of the contaminated farmland, dried through a 5mm sieve, removed plant residues and gravel, fully mixed and loaded into a 25cm×30cm plastic basin, with 8kg soil per basin. A total of 7 different treatments were set up in the experiment, including the control group CK without adding any passivator materials. Among them, attapulgite (T1), diatomite (T2) and humic acid (T3) were added with a single passivator material according to 3 % of each pot of soil weight, respectively. T4 (attapulgite + diatomite), T5 (attapulgite + humic acid) and T6 (diatomite + humic acid) were added with passivator materials according to 1.5 % of each pot of soil sample weight. Three replicates were set for each treatment. Potatoes were sown after a period of time after the application of passivator materials.

### Preparation and detection of samples

The soil samples were naturally dried in the room, and the plant residues, debris and other debris were removed and ground. The samples were passed through 10 mesh, 60 mesh and 100 mesh nylon sieves, respectively. The samples were ground and loaded into sealed bags for labeling. The whole potato plant was collected, and the surface soil was washed with tap water, and then the potato plant was repeatedly washed with deionized water for 3 ∼ 5 times, and the surface water sample was dried with absorbent paper. The potato plant was divided into roots, stems, leaves and tubers, which were put into kraft paper bags respectively. The kraft paper bags were killed at 105 ∼ 110°Cfor 30 min, and dried to constant weight at 75°C.After cooling, the stainless steel high-speed grinder was ground into powder and sifted (100 mesh) for preservation. Soil samples were measured for pH, organic matter, CEC, hydrolyzable nitrogen, soil catalase, urease, phosphatase, sucrase, soil available Cd, Pb content and chemical form changes. The content of heavy metals in each part of potato was determined.

## Results and analysis

### Effects of amendments on physical and chemical properties of heavy metal contaminated soil

It can be seen from Table 1 that compared with the control, the soil pH increased to varying degrees after the application of the modifier. The soil pH under the single application of the modifier increased by 0.27,0.14 and 0.25 units respectively compared with the control group. In the combined treatment, the combination of attapulgite + humic acid (T5) had the most obvious effect on the improvement of soil pH, which increased the soil pH to 6.18. The application of modifiers can increase the content of soil organic matter to a certain extent. In Table 1, several modifier treatments can increase the content of soil organic matter, which is 71.41 % -45.35 % higher than that of the control group. In the treatment of single modifier and combined modifier, the effect of single application of humic acid (T3) and attapulgite + humic acid (T5) is better, which is 5.41 g / kg and 10.0 g / kg higher than that of the control group, respectively.Soil cation exchange capacity is an important basis for evaluating soil fertility, improving soil and rational fertilization. From Table 1, it can be seen that the application of modifiers can effectively increase the soil cation exchange capacity. Compared with the control, each treatment increased by 4.40 cmol / kg, 2.90 cmol / kg, 3.93 cmol / kg, 6.30 cmol / kg, 7.28 cmol / kg and 3.38 cmol / kg, respectively. The application of modifiers has a significant effect on soil hydrolyzed nitrogen and available phosphorus. Compared with the control group, the content of hydrolyzed nitrogen in each treatment increased by 4.09 % ∼ 31.72 %, with an average increase of 16.29 %, while the content of available phosphorus in soil decreased to varying degrees after applying different modifier materials. Among them, the treatment of attapulgite + humic acid (T5) and attapulgite + diatomite (T4) decreased significantly, and the available phosphorus content decreased by 45.39 % and 37.77 %, respectively.

**Table 1.** Effects of amendments on physical and chemical properties of heavy metal contaminated soil.

| Treatment | pH | Organic matter<br>(g/kg) | CEC<br>(cmol/kg) | Hydrolyzable N<br>(mg/kg) | Available P<br>(mg/kg) |
| --- | --- | --- | --- | --- | --- |
| CK | 5.61±0.09 c | 18.21±0.01 d | 10.13±0.03 d | 158.67±0.15 e | 14.96±0.28 a |
| T1 | 5.88±0.03 b | 22.61±0.08 bc | 14.53±0.03 b | 185.51±0.04 bc | 12.81±0.21 bc |
| T2 | 5.75±0.04 b | 22.05±0.05 c | 13.06±0.08 c | 165.17±0.42 de | 14.02±0.12 ab |
| T3 | 5.86±0.07 b | 23.52±0.03 b | 14.06±0.08 bc | 181.17±0.81 bc | 12.13±0.12 c |
| T4 | 6.09±0.01 a | 27.77±0.02 a | 16.43±0.03 a | 192.33±0.76 b | 9.31±0.11 d |
| T5 | 6.18±0.01 a | 28.21±0.02 a | 17.31±0.05 a | 209.01±0.51 a | 8.17±0.11 d |
| T6 | 5.78±0.08 b | 22.73±0.06 bc | 13.51±0.02 bc | 173.91±0.64 cd | 13.21±0.13 bc |

Note : In the same column of the table, the same letters indicate no significant difference between treatments, and different letters indicate significant difference ( $p < 0.05$ ), the same below.

### Effects of amendments on enzyme activities in heavy metal contaminated soil and potato

It can be seen from Fig.1A that the soil urease activity in the control group was the lowest, only 0.31 mg·g·24h^-1^. After the amendment was applied, the soil urease activity was significantly increased. Compared with the control, the soil urease under each treatment increased by 0.71 mg·g·24h^-1^, 0.53 mg·g·24h^-1^, 0.67 mg·g·24h^-1^, 0.83 mg·g·24h^-1^, 0.87 mg·g·24h^-1^ and 0.55 mg·g·24h^-1^, respectively. The treatment effect of attapulgite+humic acid and attapulgite+ diatomite was the best, which was 3.81 times and 3.67 times that of the control, respectively. Fig. 1B reflects the changes of soil catalase activity after applying amendments. From the diagram, it can be seen that the application of modifiers can increase the activity of soil catalase to varying degrees. Compared with the control, the increase of soil catalase under T2 treatment is the smallest, 0.75 mL·g^-1^·h^-1^, 0.29 mL·g^-1^·h^-1^ higher than the control, 1.64 times that of the control group, and the increase of soil catalase activity under T5 treatment is the largest (1.19), 2.59 times that of the control group. As shown in Figure 1C, the activity of acid phosphatase in the soil was restored after treatment with several modifiers. The increase of soil acid phosphatase activity under the addition of each modifier was 1.28, 1.15, 1.29, 1.37, 1.48 and 1.17 times that of the control group, respectively.

**Fig. 1.**
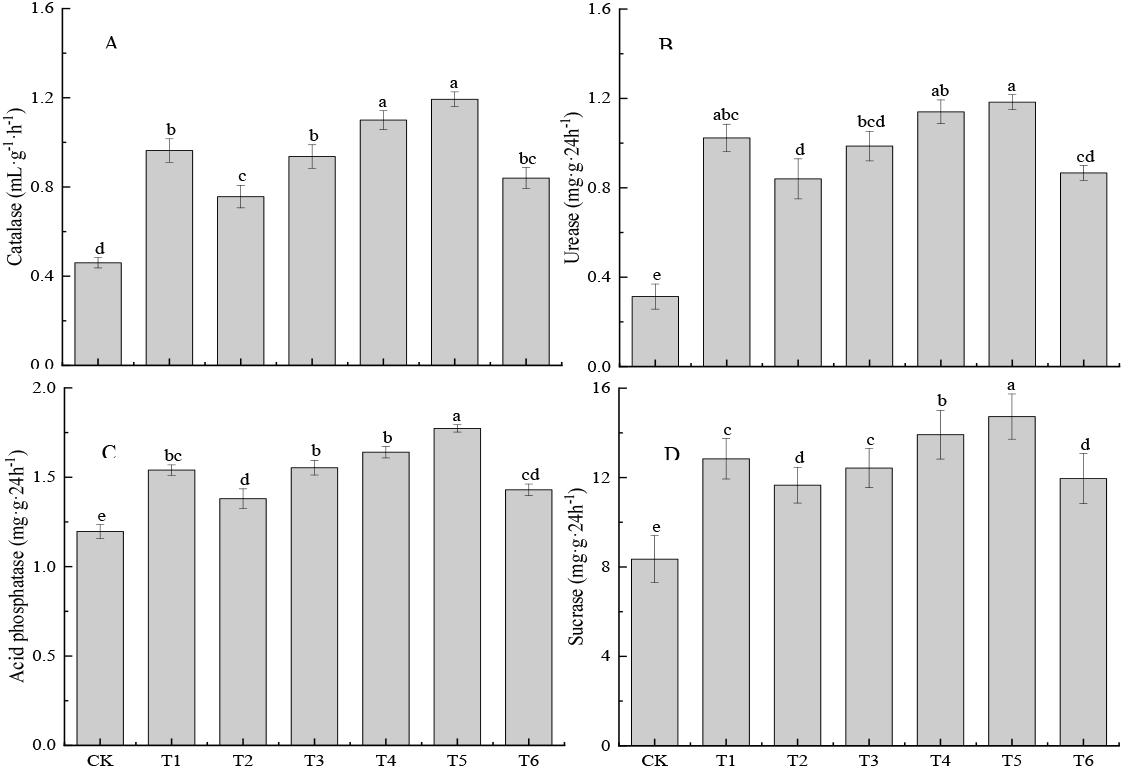
Effects of different amendments on enzyme activities in heavy metal contaminated soil.

### Effects of amendments on the available content of Cd and Pb in heavy metal contaminated soil

Fig.2 shows the changes of available contents of Cd and Pb in heavy metal contaminated soil after application of different amendments. It can be seen from the figure that the application of amendments can affect the content of available heavy metals in soil to varying degrees. Under each treatment, the content of available heavy metals in soil decreased to varying degrees, among which the content of available Cd decreased by 20.71 % -37.63 %, and the content of available Pb decreased by 12.89 % -40.95 %. Compared with the control group, the available content of Cd in soil under T5 treatment decreased to 1.4 mg/kg, a decrease of 37.63 %, and the available content of Pb decreased by 9.05 mg/kg, a decrease of 40.95 %. The available content of heavy metals in soil under T2 treatment decreased slightly. The available content of Cd in soil under this treatment decreased by 0.46 mg/kg, and the available content of Pb decreased by 2.85 mg/kg. The available content of heavy metals in soil decreased differently under different amendments, and there were differences among treatments. The overall effect was Cd : T5>T4>T1>T3>T6>T2>CK, Pb: T5>T4>T3>T1>T6>T2>CK.

**Fig. 2.**
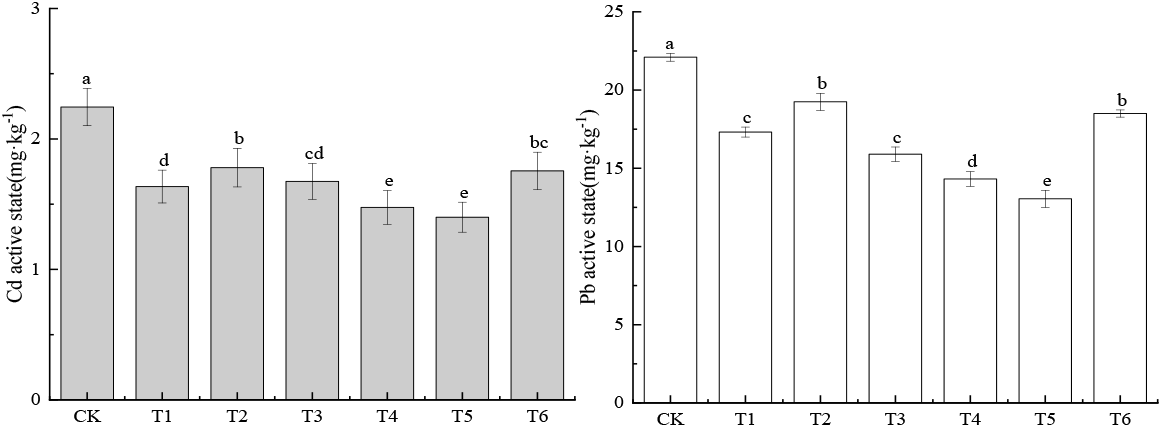
Effects of different amendments on the available content of Cd and Pb in heavy metal contaminated soil.

**Fig. 3.**
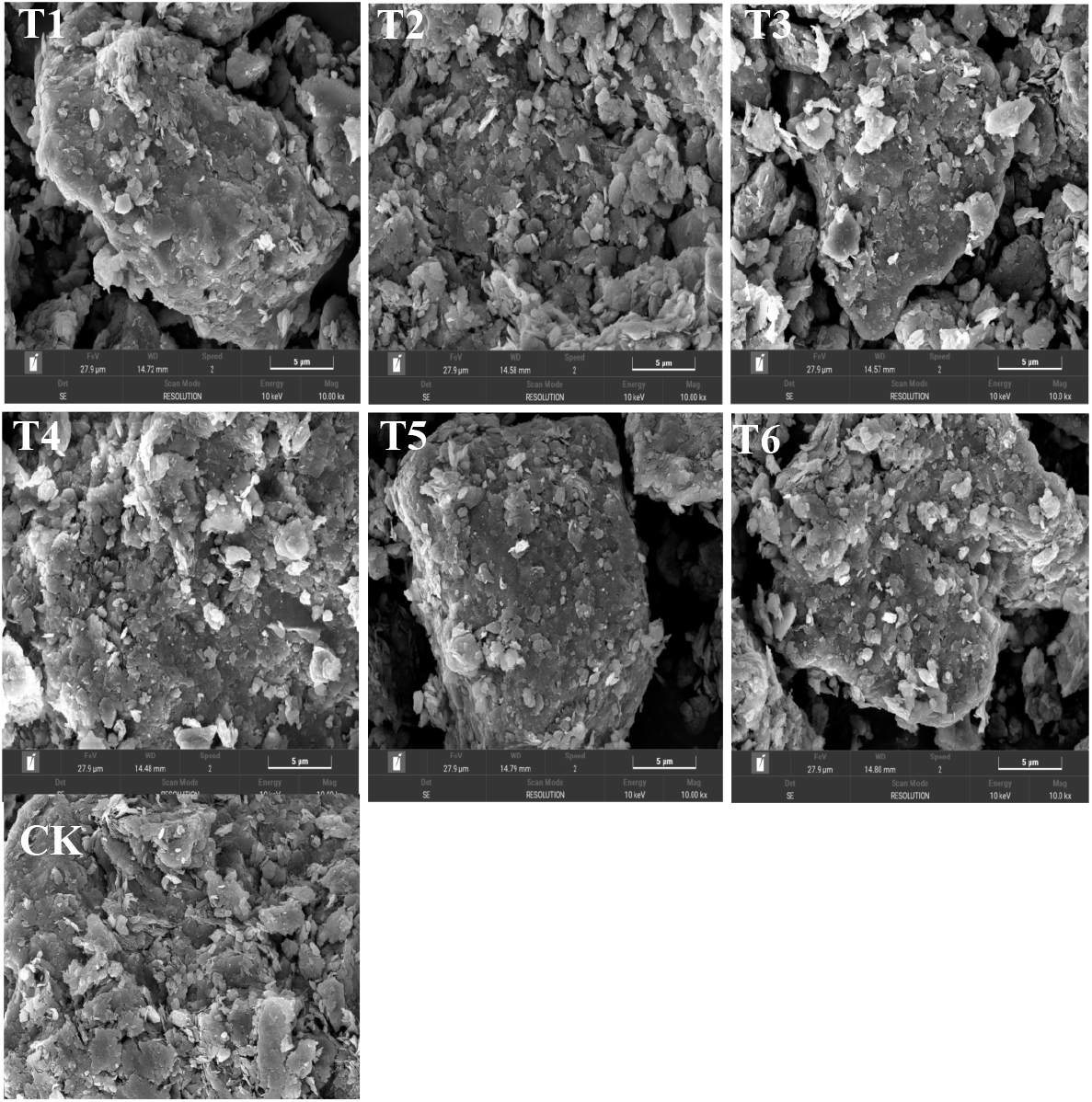
The change of soil SEM after the application of modifier.

### SEM analysis

Fig.4 shows the SEM images of soil treated with different modifier combinations. From the pictures, it can be seen that the surface morphology of soil treated with different modifier materials has changed significantly. Under CK treatment, the soil surface is irregular flake and flocculent particles. At this time, the soil structure is relatively dense, not loose, the soil pores are few, the distribution is uneven, and the heavy metal adsorption sites are sparsely dispersed. Compared with CK, after adding modifier treatment, with the increase of organic matter in the passivator, the soil surface gradually forms agglomerated flocs, the particle size increases, the voids between the aggregates also increase, the soil pore structure is more developed, and the soil permeability is improved. In addition, the increase of soil aggregate flocs also indicated that heavy metals were adsorbed on the surface after the amendment was applied to the soil to form precipitates ^**Error! Reference source not found**.^.

**Fig. 4.**
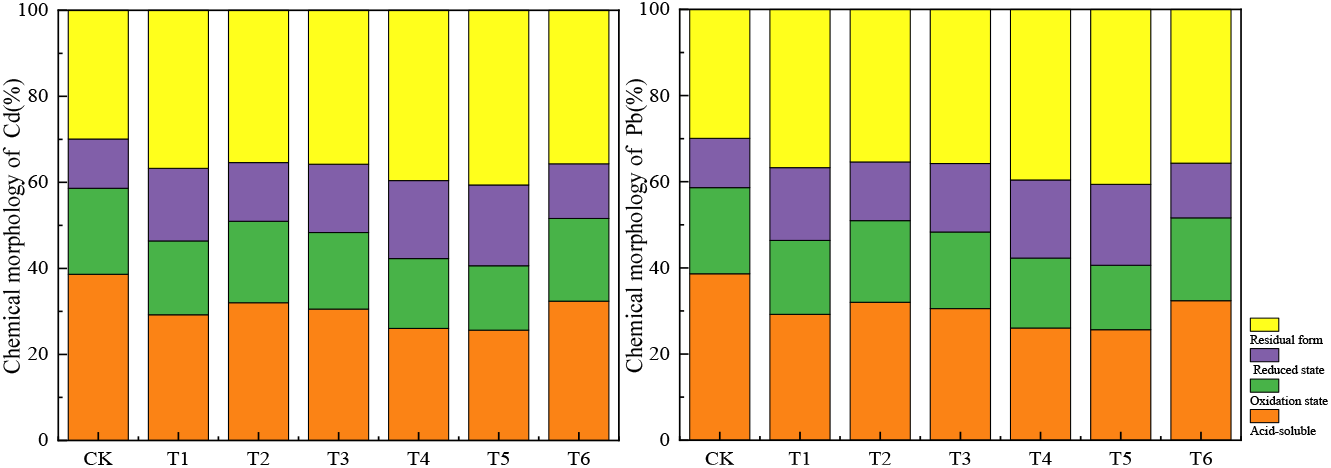
Effects of different amendments on the chemical forms of Cd and Pb in heavy metal contaminated soil.

### Effects of different amendments on the forms of Cd and Pb in heavy metal contaminated soil

Heavy metals exist in a variety of forms in the soil, which are exchangeable, carbonate-bound, iron-manganese oxide-bound, organic matter-bound and residual. Studies have shown that the total amount of heavy metals is not a good evaluation of its harm to plants and the environment. It is necessary to consider its impact after entering the soil. The speciation of heavy metals can better reflect the bioavailability and mobility of heavy metals in soil than the total amount of metals. Therefore, for the study of heavy metals, a combination of two chemical speciation classification methods can be used.The changes of chemical forms of Cd and Pb in heavy metal contaminated soil under different amendments are shown in Fig.4. From the diagram, it can be seen that the acid soluble content of Cd in the soil of CK group is 39%, the reduction state content is 20%, the oxidation state content is 11 %, the residual state content is 30%, and the percentage of the four forms of Pb is 41%, 12%, 9% and 39% respectively. After applying different amendments to the soil, the chemical forms of heavy metals in the soil changed. The acid-soluble and reduced contents of Cd and Pb in the soil under each treatment were generally lower than those of the control CK, and the acid-soluble and reduced contents of Cd and Pb in the soil with different amendments were applied. The degree of decline is different. Compared with CK, the percentage of acid-soluble Cd in soil decreased by 10%, 7%, 8%, 13%, 13% and 7%, respectively, and Pb decreased by 8 %, 4%, 7 %, 15%, 17% and 4%, respectively;compared with CK, the reduced content of Cd and Pb in soil decreased by 1%∼5% and 1%∼3%, respectively, and there was no significant difference in the reduced content of Cd and Pb among the treatments. After the amendment was applied to the soil, the oxidation state content of heavy metals Cd and Pb was higher than that of the control CK, which was different due to different treatments. Compared with CK, the oxidation state content of Cd increased by 6%, 3%, 5%, 7%, 8% and 2%, respectively, while the oxidation state content of Pb increased by 3%, 1%, 2%, 3%, 4% and 1%, respectively, compared with CK. The residual content of heavy metals Cd and Pb in soil increased with the application of amendments to varying degrees, which were higher than CK.Among them, the residual content of Cd increased by 7%, 5%, 6%, 10%, 11% and 6%, respectively, compared with the control. The residual state of Pb increased by 5%, 3%, 5%, 12%, 15% and 3%.

### The changes of heavy metal Cd and Pb contents in different parts of potato under different treatments of modifiers

Fig.5 shows the changes of Cd and Pb contents in different parts of potato under different treatments. It can be seen from the diagram that the heavy metal content in different parts of potato plants decreased under different treatments. Compared with CK, the Cd content in potato roots, stems, leaves and tubers under six treatments decreased by 41.18%∼56.61%, 45.29%∼ 58.66%, 25.90% ∼ 46.71% and 52.42% ∼ 83.50%, respectively, and the Pb content decreased to 24.30% ∼ 35.29%, 12.91% ∼26.25%, 24.15% ∼38.54% and 42.24% ∼ 55.54%. In the change of heavy metal content in potato roots, the maximum and minimum decrease of heavy metal content under different modifier treatments were Cd : T5,1.97mg/kg, 56.61%, respectively. T2,1.43mg/kg, 41.18 % ; pb : T5,3.30mg/ kg, 35.29 % ;T2,2.27mg/kg, 24.30 % ; in the changes of stems and leaves, compared with the control, the Cd content of each treatment was significantly reduced by 25.90 % ∼ 58.66 %, among which T5 decreased the most, 1.93 mg/kg and 1.1 mg/kg, respectively, compared with the control. The change trend of Pb content in stems and leaves was similar to that of Cd. The Pb content decreased the most under T5 treatment, which was 3.54 mg/kg and 2.52 mg/kg, which was 1.26 mg/kg and 1.58 mg/kg lower than that of the control group. It can be seen from the figure that compared with the control group, the Cd content in potato tubers under each treatment was significantly reduced. Among them, T5 treatment was the most significant, which was 0.43 mg/kg lower than the control, with a decrease of 83.50 %, while the Cd content in potato tubers under other treatments decreased, with a decrease of T1: 66.80%, T2: 52.43%, T3: 64.08 %, T4: 75.73% and T6: 60.19%, respectively, the content of Pb in potato tubers was different due to different modifier treatments. Compared with the control group, the content of Pb in tubers under each treatment was reduced, which was reduced by 1.54 mg/kg, 1.34 mg/kg, 1.52 mg/kg, 1.65 mg/kg, 1.76 mg/kg and 1.42 mg/kg compared with CK, respectively. There was little difference between the treatments.

**Fig. 5.**
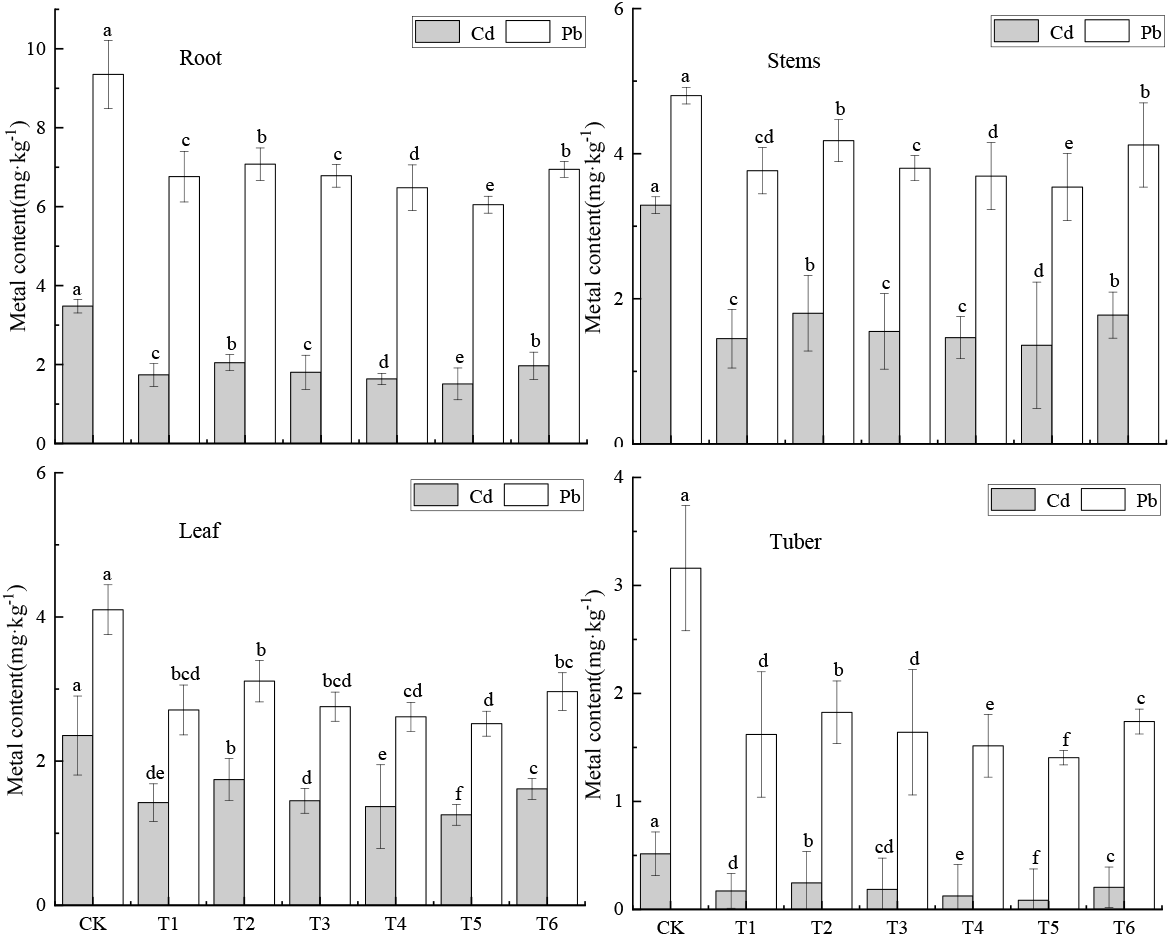
Changes of Cd and Pb contents in different parts of potato under different treatments of modifiers.

### The changes of heavy metal transport and enrichment in potato under different treatments of modifiers

Enrichment transfer coefficient can be used to evaluate the ability of plants to absorb and accumulate heavy metals. In this study, the transfer coefficient and bioconcentration factor of each part of potato were analyzed. The results are shown in Table 2. It can be seen from Table 2 that the enrichment of Cd in potato tubers was different under different modifier treatments. Compared with CK, each treatment could effectively reduce the enrichment coefficient of Cd in potato. Among them, T5 had the largest decrease, which was 73.9% lower than that of CK, and T2 had the smallest decrease, which was 39.1%. The bioconcentration factors of Cd under the other treatment combinations decreased by 56.5%, 52.2%, 65.2% and 47.8%, respectively, and the overall performance was T5<T4<T1<T3<T6<T2<CK. The enrichment of Pb in each treatment was significantly lower than that of CK (P<0.05), but there was no significant difference among the treatments. It shows that the absorption capacity of potato tubers to Cd and Pb is different under different modifier treatments. The transport coefficients of Cd in different parts of potato were also different under different modifier treatments, which were 0.06∼0.15 (root-tuber), respectively. (root-stem) 0.83∼0.95 ; (stem-leaf) 0.72∼0.98 ; The transport of Pb was (root-tuber) 0.21∼0.26 ; (root-stem) 0.51∼0.59 ; (stem-leaf) 0.71∼0.74 ; Among all treatments, T5 treatment had the greatest effect on the transfer coefficient of heavy metals in potato. The transfer coefficient of Cd in root-tuber and root-stem systems decreased by 60.0% and 9.47%, respectively, compared with CK, and Pb was 38.2% and-15.7%, respectively. In the stem-leaf transport system, the transport coefficient of Cd under each treatment was higher than that of CK, and the transport coefficient of Cd in the stem-leaf system under T5 treatment increased by 34.7%. The change of Pb in the stem-leaf system was different from that of Cd. The transfer coefficient of Pb in the stem-leaf system under each treatment was lower than that of CK, and the T5 treatment was more obvious, which decreased by 16.5%.

**Table 2.** Changes of heavy metal transport and enrichment in potato under different treatments of modifiers.

| Metal | Treatment | TF<sub>Tuber-Root</sub> | TF<sub>Stems-Root</sub> | TF<sub>Leaf-Stems</sub> | BCF<sub>Tuber-soil</sub> |
| --- | --- | --- | --- | --- | --- |
| Cd | CK | 0.15±0.05 a | 0.95±0.03 a | 0.72±0.02 c | 0.23±0.01 a |
|  | T1 | 0.10±0.01 c | 0.83±0.04 c | 0.98±0.01 a | 0.10±0.01 d |
|  | T2 | 0.12±0.01 b | 0.88±0.01 bc | 0.97±0.03 a | 0.14±0.01 b |
|  | T3 | 0.10±0.01 c | 0.86±0.01 bc | 0.94±0.02 ab | 0.11±0.01 cd |
|  | T4 | 0.08±0.03 d | 0.91±0.01 ab | 0.94±0.01 ab | 0.08±0.01 e |
|  | T5 | 0.06±0.03 e | 0.86±0.02 bc | 0.97±0.01 a | 0.06±0.01 f |
|  | T6 | 0.11±0.03 c | 0.90±0.01 ab | 0.91±0.01 b | 0.12±0.01 c |
| Pb | CK | 0.34±0.01 a | 0.51±0.05 d | 0.85±0.07 a | 0.14±0.03 a |
|  | T1 | 0.24±0.01 d | 0.56±0.08 c | 0.72±0.02 b | 0.10±0.03 b |
|  | T2 | 0.26±0.01 b | 0.59±0.06 a | 0.74±0.01 b | 0.10±0.03 b |
|  | T3 | 0.24±0.01 d | 0.56±0.01 c | 0.73±0.01 b | 0.10±0.01 b |
|  | T4 | 0.23±0.01 e | 0.57±0.05 bc | 0.71±0.01 b | 0.09±0.03 b |
|  | T5 | 0.21±0.01 e | 0.59±0.08 ab | 0.71±0.01 b | 0.09±0.05 b |
|  | T6 | 0.25±0.01 c | 0.59±0.01 a | 0.72±0.01 b | 0.09±0.03 b |

## Discuss

The absorption of heavy metals by plants is affected by many aspects, among which pH and organic matter have the greatest influence. Soil pH is a key factor affecting the activity of heavy metals in soil ^**Error! Reference source not found**.^. By increasing the concentration of OH-in the soil solution, it binds to heavy metal cations and generates insoluble precipitates, thus affecting the content of available heavy metals in the soil, while the organic matter affects the content of available heavy metals in the soil by complexing or chelating with heavy metals in the soil through the decomposition of humic acid^**Error! Reference source not found**.^. As one of the important indicators for judging soil fertility, cation exchange capacity represents the amount of soil nutrients and is an important basis for improving soil. When the pH value of the medium decreases, the negative charge on the surface of the soil colloidal particles also decreases, and the cation exchange capacity also decreases. On the contrary, it increases. In this study, the soil pH, organic matter content and soil cation exchange capacity increased after the application of different amendments (Table 1). Under the influence of soil pH and organic matter, the content of available heavy metals in soil showed a downward trend (Fig.2). Among them, T5 (attapulgite+humic acid) treatment was more obvious, which was consistent with previous studies^**Error! Reference source not found**.^. Nitrogen and phosphorus in soil solution are the main nitrogen and phosphorus sources of plants, and also an important index to measure soil fertility ^**Error! Reference source not found**.^. It has been found that clay minerals can adsorb phosphorus in soil solution and purify phosphorus through chemical mechanism, so that it can be converted into insoluble phosphorus salt to reduce the content of available phosphorus in soil solution. In this study, the content of hydrolyzable nitrogen in soil treated with different amendments was higher than that of CK, while the content of available phosphorus was lower than that of CK (Table 1), which was consistent with the results of Han**Error! Reference source not found**.

Soil enzyme can maintain the circulation of carbon, nitrogen, phosphorus and other elements in soil and maintain the biochemical balance in soil. It is an important index of soil fertility and can also characterize the pollution of soil^**Error! Reference source not found**.^. The activities of soil sucrase, catalase, urease and phosphatase directly affect the intensity of soil C, N and P biochemical cycles^**Error! Reference source not found**.^.The addition of exogenous amendments can change soil structure, moisture, temperature, salinity and nutrient concentration, and affect soil enzyme activity^**Error! Reference source not found**.^. Xu et al^**Error! Reference source not found**.^ found that the activities of soil urease, phosphatase and catalase after adding organic amendments were significantly higher than those of the control group. In this study, the activities of soil urease, catalase, phosphatase and sucrase treated with different modifiers were higher than those of the control group, indicating that the soil enzyme activity was restored after the application of modifiers, which was consistent with the conclusions of the previous studies. Among the treatments, T5 treatment had the best recovery effect on soil enzyme activity.

The ability of plants to absorb and accumulate heavy metals is expressed by bioconcentration and transport coefficients. In this study, the content of heavy metals in different parts of potato was lower than that of CK after treatment with different modifiers (Fig.5). In (Table 2), the bioaccumulation and transport coefficients of different heavy metals were different after treatment with modifiers. Under each treatment, the transport coefficient of Cd in the root-tuber and root-stem systems decreased, and increased in the stem-leaf system compared with the control, while the transport of Pb in the root-tuber and stem-leaf systems decreased, and increased in the root-stem system. The enrichment ability of potato tubers to heavy metals Cd and Pb decreased after applying modifiers. The reason may be that the application of amendments affects the chemical form and activity of heavy metals in soil solution, reduces the bioavailability of heavy metals, and thus reduces the absorption and enrichment of heavy metals by potatoes.

The change of chemical forms of heavy metals in soil will directly affect the accumulation characteristics of crops. The chemical forms of heavy metals in soil can better reflect the situation of heavy metal contaminated soil than the total amount of heavy metals in soil, and can better reflect the migration level and bioavailability of heavy metals in soil^**Error! Reference source not found**.^. Studies have shown that palygorskite combined with organic matter passivation agent can promote the transformation of Cd, Zn, Cu, Ni, Cr and Pb in heavy metal contaminated soil from acid soluble and reducing state to oxidation state and residual state compared with single passivation agent ; Natural soil passivators such as organic matter can affect the mobility of heavy metals through adsorption, complexation and oxidation reactions, and can combine organic matter with heavy metals through complexation leaching to reduce the water-soluble and acid-soluble states of heavy metals in contaminated soils ^**Error! Reference source not found**.^. Yang et al^**Error! Reference source not found**.^found that the exchangeable Cd content in soil decreased by 57% compared with the control when organic cow dung and inorganic sepiolite were applied in Cd-contaminated paddy fields. Zhang Jingjing et al added bentonite and lignite to lead-contaminated soil in different proportions, and found that Pb in contaminated soil was transformed from acid-soluble state to residual state after ion exchange, hydrolysis and adsorption of bentonite and chelating adsorption of humic acid in lignite. It can be seen that soil amendments of different materials can have a positive impact on the chemical forms of heavy metals in soil, thereby reducing the absorption and accumulation of heavy metals by plants and reducing the stress of heavy metals on plants. In this study, after applying different amendments, the acid-soluble and reduced states of soil heavy metals Cd and Pb can be transformed into oxidized and residual states, which is consistent with the previous research conclusions.

## Conclusion

After applying different modifiers, the pH, organic matter and cation exchange capacity of heavy metal contaminated soil can be improved to varying degrees. At the same time, it can also affect the change of soil nitrogen and phosphorus content and improve soil physical and chemical properties.

After the application of amendments, soil urease, catalase, acid phosphatase and sucrase were improved to varying degrees, indicating that the amendments could have a positive effect on the activity of some enzymes in the soil, and the metabolic reactions of some enzymes in the soil could be restored.

The application of amendments can reduce the content of DTPA-extractable Cd and Pb in soil, cause the change of chemical forms of Cd and Pb, reduce the activity of heavy metals in soil, significantly reduce the content of Cd and Pb in potato plants, and reduce the enrichment and transport of Cd and Pb in potato plants. Among them, the effect of amendments under T5 (attapulgite + humic acid) treatment is better.

## Declare

### Availability of data and materials

All data generated or analyzed during this study are included in this published article.

### Funding

This work was supported by the Provincial Science and Technology Plan of Guizhou,China(20203001)and the National Natural Science Foundation of China (No 21866008)

### Author contribution

In this study, Dr. Liu Ke provided the guidance of theoretical knowledge, including the adjustment of the rationality of experimental design and so on. Mr. Zhao Changchun is the practitioner of the research, including the implementation of the whole experiment and the summary of the results. Ms. Long Li, Ms Lili Zhang and Mr Shiqing Feng provided help during the experiment.

